# Gene-Culture Coevolution Favours the Emergence of Traditions in Mate Choice through Conformist Social Learning

**DOI:** 10.1101/2025.09.05.674569

**Authors:** D. Federico, M. Cosme, F.X. Dechaume-Moncharmont, J.B. Ferdy, A. Pocheville

## Abstract

The emergence of cultural traditions has long been considered to depend on sophisticated social learning mechanisms, particularly conformist transmission exhibiting a disproportionate bias towards majority behaviours. We challenge this assumption by demonstrating that gene-culture coevolutionary dynamics can fundamentally alter the conditions required for tradition formation, especially in the context of mate choice. Using simulation models of mate choice where female preferences are socially transmitted and male traits are genetically inherited, we show that the interaction between cultural and genetic transmission creates novel evolutionary dynamics. In a first model (the ‘Cultural Model’) representing classical cultural evolution scenarios, we show that traditions emerge only when conformity disproportionately amplifies majority behaviours. In a second model (the ‘Gene-Culture Model’), we introduce a distinction between learned behaviours and realized behaviours and show that when mate choice depends on both conformist learning and male trait availability, even simple social learning strategies can establish persistent traditions. In this context, majority exaggeration is no longer a necessary property of social learning for traditions to arise. This occurs because positive frequency-dependent interactions between preference and trait dynamics provide an additional mechanism that reinforces majority behaviours beyond conformist learning. Our findings have broader implications for cultural evolution theory, suggesting that many behaviours that depend on resource availability may exhibit tradition-like patterns under less stringent social learning conditions than previously assumed. By distinguishing between preference and choice, and demonstrating the importance of gene-culture interactions, this study advocates for more integrated approaches to understanding cultural evolution that consider the interplay between multiple inheritance systems.

## 1 Introduction

Cultural evolution, long recognized exclusively in humans, is now studied in numerous other animal species [1, 2]. Among the most extensively investigated examples are the cultural transmission of songs in various birds and cetaceans [3, 4, 5, 6, 7], as well as the cultural transmission of food acquisition techniques among several primate species [8, 9]. The number of candidate contexts and species has been growing rapidly in the last decade (see recent reviews on culture in mammals [10], cetaceans [11], monkeys [9], great apes [12], birds [13], fish [14] and insects [15]). While this diversification of species studied through the lens of cultural evolution has led to a refinement of theoretical questions tailored to specific models, it also raises the broader issue of identifying the simplest or most fundamental forms of cultural evolution – and the challenge of detecting them.

A fundamental pattern of cultural evolution at the population level appears to be the presence of traditions [16]. A broad definition of tradition was suggested by Fragaszy and Perry [17] as ‘a distinctive behavior pattern shared by two or more individuals in a social unit, which persists over time and that new practitioners acquire in part through socially aided learning’ – that is, through the acquisition of knowledge from conspecifics. Traditions, which have come to be a defining feature of culture in many studies [18, 19, 20], refer in other words to any behaviour that: (1) is transmitted at least in part through social learning, (2) is sufficiently widespread within the population, and (3) exhibits a degree of temporal stability – that is, persists across multiple generations. One challenge in the study of cultural evolution implies to better understand how traditions emerge, spread, persist, or disappear over time. Traditions have thus far been studied primarily through the lens of social learning [21] – understandably, as it is a fundamental and necessary mechanism in cultural evolution. However, social learning is not the only driver of cultural dynamics and a growing body of research highlights the importance of considering how social transmission interacts with other factors – such as the population age structure [22], the social network configuration [23], or the dynamics of dispersal [19] – to better understand cultural evolution [24, 21]. Several authors have also stressed the value of studying cultural and genetic evolution jointly, as their interaction can generate complex gene–culture coevolutionary dynamics [25, 26, 27]. Theoretical modelling is essential for understanding how cultural patterns emerge in these more complex evolutionary settings, and for disentangling the respective roles of social learning and other processes in the formation of traditions.

Although it has received less attention than other behaviours in cultural evolution studies, mate choice provides a promising context for investigating the emergence of traditions [18, 28]. This is because mate choice combines empirical evidence for social learning with theoretical predictions of frequency-dependent dynamics that could favour traditions. Consistent with this idea, a substantial body of empirical research highlights the role of social learning in shaping mate choice across a wide range of species [29, 30, 31]. In particular, the phenomenon of ‘mate-choice copying’ – where individuals replicate the mate choices of others – has been well documented [32, 33, 34]. Moreover, conformist social learning (hereafter referred to as conformity), which involves copying the majority, has been demonstrated in mate choice contexts in species as diverse as humans and fruit flies [35, 18]. Originally defined by Boyd and Richerson as the ‘disproportionate probability of adopting the most common variant’ [36], conformist transmission is a form of social learning that is widely regarded in cultural evolution studies as a key mechanism promoting the emergence of traditions [37, 38, 39]. The term ‘disproportionate’ implies that when a given variant is exhibited by, for example, 70% of individuals in the group, a conformist learner will adopt it with a probability greater than 70%. By amplifying and reinforcing majority behaviours, conformity is usually expected to increase within-group homogeneity while fostering between-group divergence – two hallmark features of traditions [28]. In fruit flies, observer females were found to significantly copy the majority choice among a group of 5 or 6 demonstrator females – each observed mating with one of two coloured male types [18]. Observers copied the most frequent choice regardless of the strength of the majority (i.e., 3/5, 4/6, 5/6, or 6/6), and a model extrapolating these experimental results to the population and multi-generational scale predicts the possibility of long-term traditions in mate choice [18].

Beyond empirical evidence for social learning and conformity, another reason to consider mate choice as a promising context for studying traditions relates to the predicted runaway dynamics in certain sexual selection scenarios. Classical models of sexual selection have shown that mating preferences in one sex can coevolve with traits under preference in the other, sometimes giving rise to reinforcing dynamics such as the Fisher runaway process [40, 41, 42]. While these models traditionally assume that both traits and preferences are genetically transmitted, more recent work has incorporated socially learned preferences and found that positive feed-back loops between the evolution of preference and trait can also exist in this context [43, 44, 45]. The frequency-dependent interactions between socially transmitted preferences and genetically inherited traits may significantly alter the conditions under which cultural traditions emerge, which has not been explored in these previous models. They have instead primarily focused on the evolutionary consequences for male trait dynamics – that is, on the genetic component only – while overlooking the implications for cultural evolution.

In this theoretical study, we explore the idea that traditions in mate choice may be shaped by gene-culture coevolution: an interplay between the social transmission of preferences and the genetic transmission of traits. We hypothesise that this interplay can modulate the standard conditions for the emergence of traditions, and we investigate this prediction through a series of simulation models. Concretely, we examine the conditions for the emergence of traditions in mate choice using simulation models in which female preferences are socially transmitted through conformity, while the male traits preferred by females are genetically inherited. In a first model, the ‘Cultural Model’, we isolate the effects of social transmission alone on the emergence of traditions. In this setting, female choices depend solely on socially acquired preferences and are unaffected by the frequency of male traits – meaning that social transmission is independent of genetic transmission. This model follows classical frameworks in cultural evolution, where social transmission is the primary evolutionary force, and it serves as a baseline for comparison with our second – and arguably more realistic – model. In this second model, the ‘Gene–Culture Model’, we introduce the possibility of gene–culture coevolution between traits and preferences by making female choices dependent on the frequency of male traits, and we explore how this coevolution influences the conditions under which traditions emerge.

Most of the cultural evolution literature adopts the definition of conformist transmission proposed by Boyd and Richerson [36, 46, 39], which requires a disproportionate probability of adopting the majority behaviour – a feature we term *majority exaggeration*. This conformist transmission is typically modelled as an S-shaped relationship between the frequency of a behaviour at time *t* and the probability of adopting this behaviour at *t* + 1, often called the *conformity curve*. This definition, based on majority exaggeration in the conformity curve, focuses on population-level outcomes of conformity – namely, the amplification of majority behaviours – rather than on underlying mechanisms. In models where conformity is the sole driver of behavioural change, majority exaggeration is necessary for traditions to arise. However, it may become unnecessary when other reinforcing processes are at play – such as those expected in sexual selection contexts. Moreover, majority exaggeration has proven difficult to demonstrate empirically, even when individuals appear to employ majority-copying strategies [47, 48]. Importantly, Boyd and Richerson’s definition of conformity focuses on realized behaviours. However, realized behaviours need not result solely from social learning: they may deviate from learned preferences when behaviour depends on external constraints, such as resource availability. As Morgan and van Leeuwen [39] note, conformity theory has not yet explored this divergence between learned and realized behaviours.

In this paper, we adopt a more mechanistic approach to modelling conformity. Rather than imposing a population-level conformity curve, we model individuals who engage in a conformist learning process, distinguishing between mate *preference* and mate *choice*. Based on two parameters reflecting individual capacities – observation capacity and copying fidelity – individuals first observe a sample of mate choices and detect which male type is most frequently chosen, thereby forming a preference for that type. They then choose a mate, whose type may differ from their preferred type due to copying errors or, in our Gene-Culture Model, due to male type availability. This approach allows us to assess which individual capacities give rise to traditions at the population level and enables conformity curves to emerge from individual behaviours rather than being imposed a priori. This enables us to examine how conformity interacts with genetic evolution, and determine whether majority exaggeration is a necessary property of social learning for tradition to emerge when preferences and traits co-evolve. Overall, we find that gene-culture interactions facilitate tradition emergence, allowing traditions to arise and persist even without majority exaggeration in social learning.

## 2 Methods and results

### 2.1 The models

We developed individual-based models using R version 4.4.0 [49] to investigate the conditions under which traditions emerge in mate choice.

#### 2.1.1 Models’ overview: the life cycle

The population consists of male and female individuals, each belonging to one of two age classes: juveniles or adults. At each time step, juveniles transition to adulthood with a probability *a* (creating overlapping generations when *a <* 1). Once they reach adulthood, individuals live for a final time step, during which they mate and reproduce. Females are the choosy sex and mate only once, whereas males can mate multiple times (up to *m* times) if chosen by different females. Juvenile females learn their mate preferences by observing the mate choices of adult females. The number of newborns produced at each time step is set to keep the population size *K* constant. The parents of each newborn are randomly sampled (with replacement) from the pool of all possible reproducing pairs. Sex is assigned at random, with an equal probability of being male or female.

#### 2.1.2 Individual traits

Each individual is characterized by a set of binary variables (0/1) that define their traits and follow distinct transmission pathways. These variables include: (1) a male trait, which is genetically inherited, carried by both sexes but expressed only in males, distinguishing two recognizable male types (type 0 and type 1, with no inherent difference in survival or mating opportunities); (2) in females only, a preference for a specific male type (0 or 1), which is socially learned; (3) in females only, a mating choice for a specific male type (0 or 1), which is the expressed behaviour based on the female’s socially learned preference and can deviate from this preference due to copying errors and/or constraints on male type availability. The core of our models lies in how these traits are transmitted across generations, following one of two transmission pathways: genetic or social. The fidelity of these transmission paths is controlled by three key parameters, summarized in table 1 and detailed below.

**Table 1:** The three parameters influencing the fidelity of genetic and social transmission in our models. Social transmission fidelity is determined by *n_obs_*and *c*, whose combination defines the intensity of conformist social learning in females. Genetic transmission fidelity is governed by parameter *h*, which affects the heritability of male traits. *n_mat_* represents the total number of matings occurring at one time step.

| Parameter | Interpretation | Range | Transmission Path |
| --- | --- | --- | --- |
| $n_{obs}$ | **Number of observations:** the finite number of adult pairings that a juvenile female can sample and observe, through which she identifies and develops a preference for the most commonly (majority) chosen male type. | $[1, n_{mat}]$ | **Social** – involved in the preference acquisition step in conformity |
| $c$ | **Copying fidelity:** the probability that an adult female successfully memorizes and applies her learned preference when selecting a mate. A value of $c = 0.5$ represents the null model, where no social transmission occurs. | $[0.5, 1]$ | **Social** – involved in the mate choice step in conformity |
| $h$ | **Genetic transmission fidelity:** controls the probability of faithful male trait transmission from parent to offspring (analogous to heritability). | $[0, 1]$ | **Genetic** |

#### 2.1.3 The path of genetic transmission: heredity of the male trait

The male trait is genetically determined and is inherited from either the mother or the father with a probability of 1/2. Parameter *h* represents the fidelity of trait transmission from parent to offspring, analogous to heritability in quantitative genetics, and ranging from *h* = 0 (no transmission) to *h* = 1 (perfect transmission). Concretely, for each newborn: with probability *h*, the male trait matches the parental trait; with probability 1 *− h*, the trait is randomly assigned (type 0 or 1 with equal probability). This modelling choice ensures that when *h* = 0, newborns’ traits are independent of parental traits, providing a null model without genetic transmission – which also allows comparison with Danchin et al.’s model [18], where male traits are not inherited.

#### 2.1.4 The path of social transmission: acquisition of preference and mate choice through conformity in females

In our models, females’ preference for a male type is exclusively socially transmitted. Social transmission follows a conformist social learning process that unfolds in two sequential steps at the individual level: during the juvenile stage, females acquire their preference from reproducing adult females, and in adulthood, they use this learned preference to choose a mate. The fidelity of social transmission is influenced by two parameters, *n_obs_* and *c* (see table 1 and below).

##### 2.1.4.1 Step 1: Preference Acquisition

Preference in juvenile females is socially acquired by observing adult females’ mate choices. At each time step, every juvenile female samples *n_obs_* adult mating events, randomly drawn from all matings occurring during that step. She then adopts a preference for the male type that appears most frequently in her sample – that is, that corresponds to the majority choice. If both male types are equally represented in the sample, the preference is randomly assigned with equal probability (figure 1, preference acquisition). There is no memory between time steps: if a female remains in the juvenile stage for multiple steps, her preference is reset at each step and re-learned based on new observations. If the number of matings is lower than *n_obs_*, females simply observe all available matings. We assumed: (1) that all females in the population share the same *n_obs_*value, and (2) that the majority behaviour within a sample can always be detected, regardless of whether the majority is slight or pronounced. While conformity requires *n_obs_ >* 2 (so that a majority can be identified), we also explored the case of *n_obs_*= 1, known as ‘single copying’ – a scenario widely used in mate-choice copying experimental studies.

**Figure 1:**
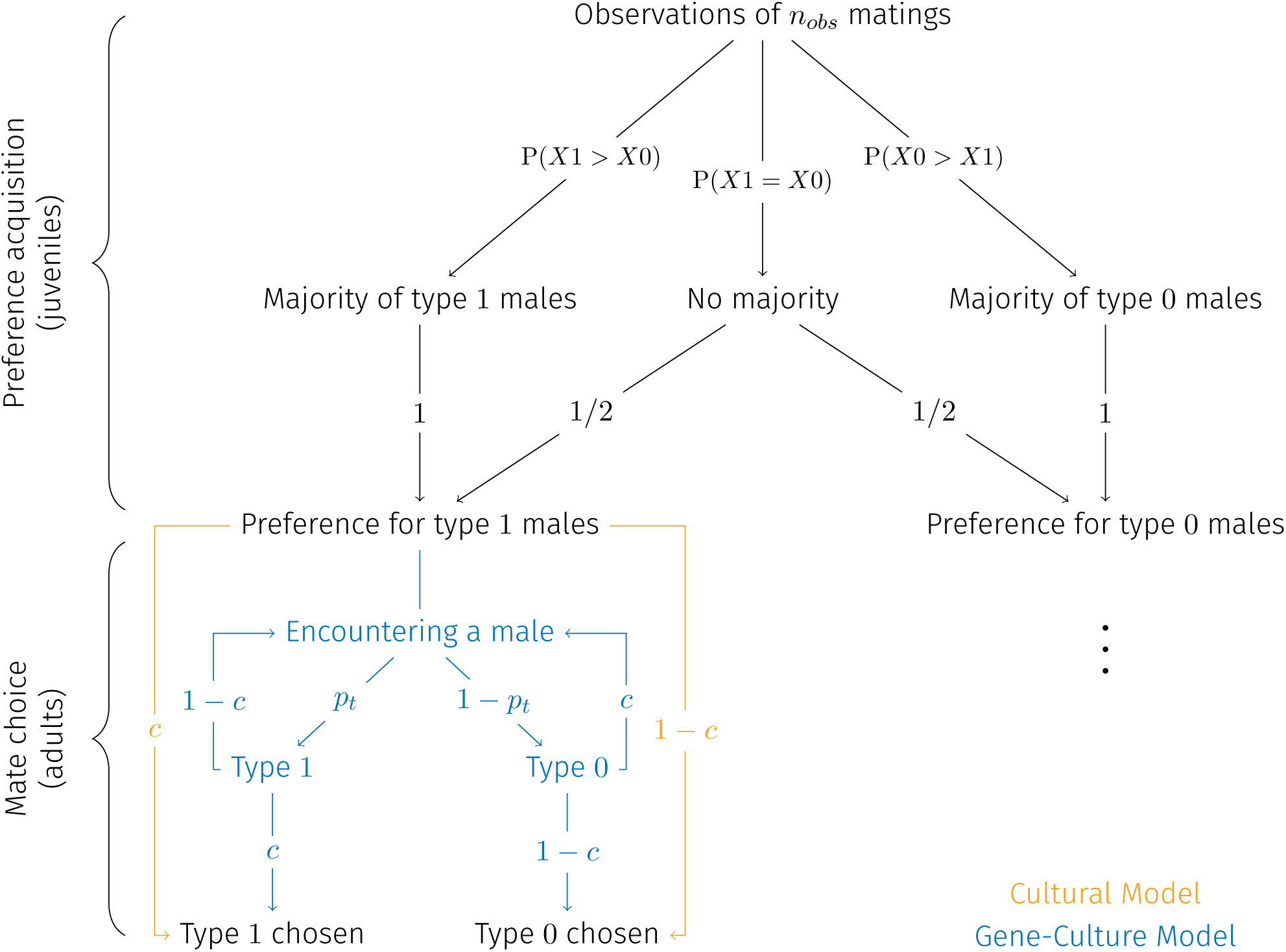
The steps of preference acquisition and mate choice through conformity in our models. Nodes represent possible events, while arrows indicate transitions between events along with their associated probabilities. The conformity process in females unfolds in two stages: i) *Preference acquisition* (juvenile stage): each female samples and observes *n_obs_* matings from the previous time step, learning to prefer the majority male type within these observations. If no majority is present, preference is assigned randomly. The probabilities to observe a majority for type 1 P(*X*1 *> X*0), for type 0 P(*X*0 *> X*1), or no majority P(*X*1 = *X*0) are mathematically developed in the section ‘Computing conformity curves’. ii) *Mate choice* (adult stage): this step differs between models: A) Cultural Model (orange): females have access to all males and choose based solely on their learned preference, modulated by their copying fidelity *c*. B) Gene-Culture Model (blue): mate choice depends not only on preference (modulated by *c*) but also on the probability of encountering each male type. This probability is equal to the relative frequency of each type among adult males (*p_t_* for type 1 males and 1 *− p_t_* for type 0 males). In the Gene-Culture Model, females randomly encounter males until they accept one. For clarity, only the mate choice process for females preferring type 1 males is illustrated, as the process is symmetrical for those preferring type 0 males.

##### 2.1.4.2 Step 2: Mate Choice

Mate choice represents the transition from a female’s learned preference to her actual choice of a mate during adulthood. To account for potential errors in conformist learning, we introduced a ‘copying fidelity’ parameter (*c*), which quantifies the probability that an adult female successfully follows her learned preference when she encounters a male of her preferred type. The implementation of mate choice and the interpretation of parameter *c* differ between the two models we examined, as described below (summarised in figure 1).

**Cultural Model** In this first model, mate choice occurs independently of male type frequencies in the population, a scenario we refer to as ‘unconstrained mate choice’. In this scenario, as long as the two types of males exist in the population, the female chooses a male of its preferred type with probability *c*, or a male of the non-preferred type with probability 1 *− c* (figure 1, mate choice in orange). This Cultural Model follows classical approaches to conformist learning in cultural evolution, where the adoption of socially transmitted behaviours is primarily driven by social learning capacities, without additional external constraints [36, 28, 50]. We use this model as a first step to explore how varying the strength of conformist social learning — modulated by *n_obs_* and *c* — affects the emergence of traditions in mate choice within a simplified context. It provides a baseline for comparison with our second, and arguably more realistic, model.

**Gene-Culture Model** In contrast, mate choice in our second model is partially constrained by the frequency of male types in the population (resource availability), a scenario we refer to as ‘constrained mate choice’. In this scenario, each female sequentially encounters males and decides for each whether she accepts to mate. More precisely, at each encounter: (1) the probability that the encountered male is of type 1 corresponds to the frequency *p_t_* of that type in the adult male population at generation *t*; (2) if the male is of the female’s preferred type, she accepts to mate with him with probability *c* (and refuses with probability 1 *− c*). If the male is not of the female’s preferred type, she accepts to mate with probability 1 *– c* and refuses with probability *c*; (3) the female continues encountering new males until she accepts one or no more males are available (which can occur when males reach their maximum number of allowed matings, *m*). With our baseline parameter *m* = 100, males never reach their mating limit in practice, but we explore the effects of lower values of *m* in Section D of Supplementary Materials. This sequence of events is represented in figure 1 (mate choice in blue). Note that females’ sequences of encounters occur within a single model time step (their adult stage). By constraining mate choice based on male type frequencies, the Gene-Culture Model introduces potential interactions between the dynamics of socially transmitted preferences and genetically inherited traits. We use this model to investigate how a potential gene-culture coevolution context influences the emergence of traditions in mate choice.

Mate choice in both models converge in the specific case of *c* = 1, where females exclusively mate with a male of their preferred type (’perfect copying’ scenario, where preference directly translates into choice). In this limiting case, females remain unmated if all available males are of the non-preferred type.

We refer to the first model as the ‘Cultural Model’ because cultural evolution occurs independently of genetic dynamics (unidirectional interaction), and to the second as the ‘Gene-Culture Model’ because it involves reciprocal interactions between genetic and cultural evolution (bidirectional interaction, i.e., coevolution). Note that both models include genetic inheritance of male traits and social transmission of female preferences; the distinction lies in whether these processes interact bidirectionally.

**Null Model: Random Mate Choice** The models also converge when *c* = 0.5, a scenario in which mate choice is entirely independent of preference, resulting in random choices with no influence from social learning. We use the *c* = 0.5 scenario as a null model for both the Cultural and Gene–Culture Models. Comparing this null model to its respective social counterparts allows us to isolate the specific effects of social transmission on evolutionary dynamics.

### 2.2 Emerging conformity curves and majority exaggeration

Conformity curves typically describe the relationship between the frequency of an observable behaviour in the population at time *t* and the probability that a conformist learner will adopt this behaviour at *t* + Δ*t*. These curves are often assumed to follow an S-shape, with the probability of adopting the majority behaviour exceeding its frequency in the population – a key characteristic of ‘conformist transmission’ as originally defined by Boyd and Richerson [36], which we term majority exaggeration. In this study, we did not impose a predefined shape for the conformity curves. Instead, we allowed them to emerge from the individual behaviours described above and examined whether these emerging conformity curves exhibited majority exaggeration in at least some region. We also ensured that what we term conformity curvesreflect only social learning by removing the effects of external constraints (male type availability) on mate choice.

#### 2.2.1 Probability vs. Conditional Probability to Choose one Type of Males

In the Gene-Culture Model, mate choice is influenced by both preference and encounters with males. To separate the two effects, we distinguished between two probabilities: the overall probability of choosing a male of type *i*, denoted as *P* (*C_i_*), and the conditional probability of choosing a male of type *i* given that such a male has been encountered, denoted as *P* (*C_i_|E_i_*). The probability *P* (*C_i_*) is influenced by females’ learning abilities (*n_obs_* and *c*) and by the frequencies of male types (*p_t_* and 1*−p_t_*). In contrast, the conditional probability *P* (*C_i_|E_i_*) depends solely on females’ learning abilities.

In principle, the conditional probability *P* (*C_i_|E_i_*) cannot be computed in the Cultural model, Observations of *n_obs_* matings where encounters are not explicitly modelled. However, the concept remains applicable by considering that females in this model encounter simultaneously the two types of males. This implies that *P* (*E_i_*) = 1, whatever the frequency of type *i* is. It then follows that, in the Cultural model, *P* (*C_i_*) = *P* (*C_i_|E_i_*).

Finally, because the conditional probability *P* (*C_i_|E_i_*) eliminates the effects of male encounters and depends only on females learning capacities, it is identical in the Cultural and Gene-Culture models.

In this work, we define the ‘conformity curve’ as the relationship between the conditional probability *P* (*C*_1_*|E*_1_)*_t_*of females choosing type 1 males at time *t*, and the frequency *F* (*C*_1_)*_t−_*_1_ of choices for type 1 males in the previous generation *t −* 1.

#### 2.2.2 Computing Conformity Curves

The conditional probability of choosing type 1 males can be computed analytically as:

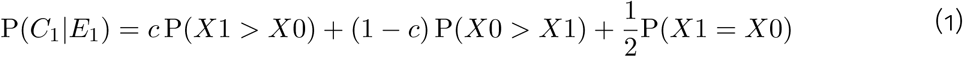

where *X*1 (respectively *X*0, with *X*1 + *X*0 = *n_obs_*) is a random variable that corresponds to the number of mating with males of type 1 (respectively type 0) a juvenile female has observed. The probabilities P(*X*1 *> X*0), P(*X*0 *> X*1) and P(*X*1 = *X*0) are thus the probabilities of observing a majority of choices for type 1, type 0, or no majority at all during the juvenile stage. As a female never observes the same mating pair twice, the distribution of *X*1 is a hypergeometric with

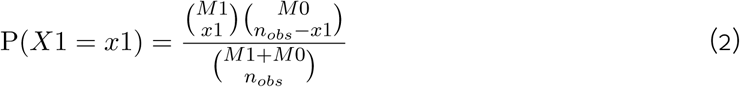

where *M*_1_ (respectively *M*_0_) is the total number of matings with type 1 (respectively type 0) males in the population, which depends on *K* and on the demographic parameters.

We used equation (2) to compute *P* (*C*_1_*|E*_1_)*_t_* for various values of of *n_obs_* and *c*, resulting in the conformity curves shown in figure 2a. We verified the validity of our analytical results by comparing the analytical curves with those obtained from simulations in both the Cultural Model and the Gene–Culture Model, and found that they match closely (see Supplementary Materials, Section A, figure S1).

**Figure 2:**
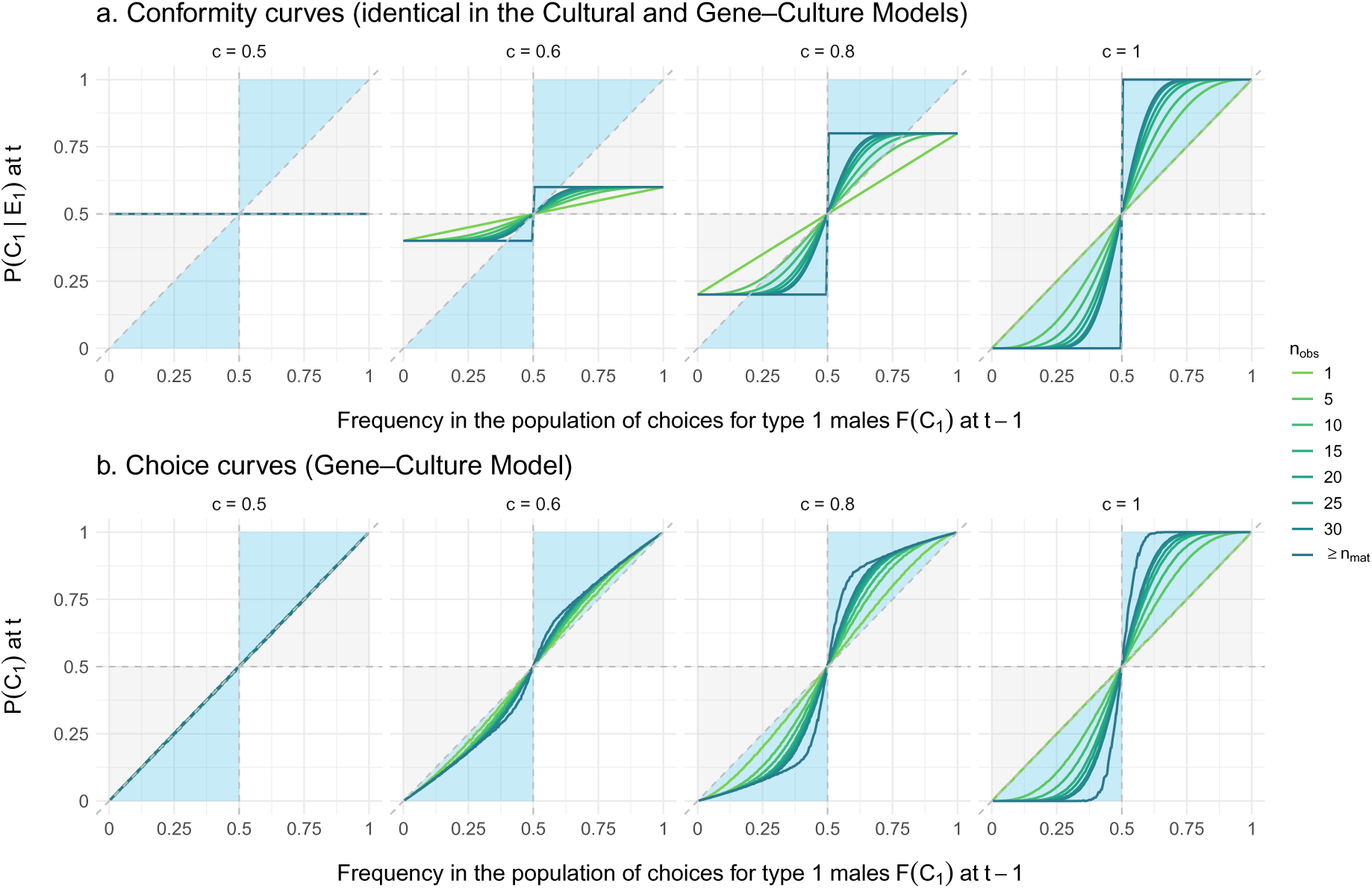
Conformity curves and choice curves emerging from our models. All curves were computed for populations of 1000 individuals. **a.** Conformity curves (Cultural and Gene-Culture models) for various combinations of *c* (different panels in each row) and *n_obs_* (different line colours, with *n_mat_* being the total number of matings occurring per time step). Conformity curves represent the relationship between the conditional probability for females to choose type 1 males at *t* (given an encounter with a type 1 male), and the frequency of choices for type 1 males in previous females at *t −* 1. Since they are independent of male type frequencies, these curves reflect only conformist learning capacities and are identical for both models. They were computed analytically. Conformity responses exhibiting majority exaggeration correspond to curves intersecting the blue regions, while those without majority exaggeration fall within the grey regions. **b.** Choice curves (Gene-Culture Model) for various combinations of *c* and *n_obs_*. Choice curves represent the relationship between the probability for females to choose type 1 males at *t*, and the frequency of choices for type 1 males in previous females at *t −* 1. Unlike conformity curves, choice curves are influenced by male type frequencies (in the Gene-Culture Model only). They were derived from 1000 simulation replicates by computing the frequency of females choosing type 1 at *t*_1_ for different fixed values of the frequency of females choosing type 1 at *t*_0_ (using the baseline parameter values described in the section about simulations).

For the Gene-Culture Model, we derived ‘choice curves’ in addition to ‘conformity curves’ and estimated *P* (*C*_1_) through 1000 replicate simulations for each value of *n_obs_* and *c* we have tested (figure 2b). Specifically, *P* (*C*_1_) was estimated as the frequency of females choosing type 1 males at generation 1 for different fixed proportions of females choosing type 1 at generation 0 (using the baseline parameter values described in the section below on simulations).

#### 2.2.3 Majority Exaggeration

Among these conformity curves, we identified those that exaggerate the majority choice in at least part of their frequency range – that is, curves where the probability of adopting the majority behaviour exceeds its frequency in the population for at least some frequency values. Formally, this occurs in regions where:

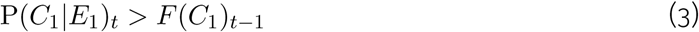

Conformity responses displaying majority exaggeration correspond to the curves that intersect the blue regions in figure 2a, whereas those without majority exaggeration fall entirely within the grey regions.

#### 2.2.4 What we Learn from Conformity Curves

These population-level curves provide a clear visualization of the range of social learning responses emerging from our models. For *c* = 0.5 (null model, without social transmission), the response remains constant, as females choose randomly. For *c >* 0.5 and *n_obs_ ≤* 2, the response is linear because females copy one or two randomly observed older females, with no majority influence involved. This encompasses what is typically referred to as single-copying. For *c >* 0.5 and 2 *< n_obs_ < n_mat_*, conformity curves take a sigmoidal shape, where *n_obs_* affects the slope at the inflection point, while *c* determines the height of the two plateaus. For *c >* 0.5 and *n_obs_* = *M* 1 + *M* 0, i.e. when females observe all the mating events in the population, the response becomes a step function, with *c* again determining the plateau heights. Importantly, these curves highlight which combinations of *n_obs_* and *c* generate majority exaggeration – that is, the curves intersecting the blue regions in figure 2a.

A key distinction in the Gene-Culture Model is that, as defined above, choice curves differ from conformity curves (compare figures 2a and 2b), whereas in the Cultural Model, they are identical. When choosing type 1 is the majority behaviour at *t −* 1, the probability of choosing type 1 at *t* is significantly higher in the Gene-Culture than in the Cultural Model. This is due to the interaction between conformity and the frequency of available type 1 males, which reinforces the probability of choosing type 1.

Our goal in the following was to investigate whether these different social learning responses, along with their potential interactions with genetic evolution, can give rise to traditions.

### 2.3 The detection of traditions

#### 2.3.1 Defining a Detection Threshold for Traditions

We analysed the temporal dynamics of choices to determine the conditions that lead to traditions. Traditions are generally described as the persistence of a socially transmitted behaviour across multiple generations [17], yet the exact number of generations required for a behaviour to qualify as a tradition is rarely specified. In this study, we propose a quantitative threshold for tradition detection by comparing the persistence of a majority behaviour in the presence (*c >* 0.5) and absence (*c* = 0.5) of social transmission.

Our method for detecting traditions proceeded in two steps. First, we defined a detection threshold using the null model (c = 0.5), where choices evolved solely under the influence of drift and mutation. To do so, we ran 100 null model simulations and, for each replicate, discarded the first 100 time steps to eliminate effects of initial conditions. We then measured the persistence duration of the majority choice observed at time step 100 (i.e., how long this choice remained the majority). The 95th percentile of these persistence durations across all null model replicates defined our detection threshold. Second, we applied this threshold to the non-null models (c > 0.5). Using the same procedure (discarding the first 100 time steps and measuring persistence from step 100), we classified any majority choice that persisted beyond the threshold as a tradition. All simulations were run for a maximum of 1,000 time steps, and only the first emerging tradition was retained in our analyses. Examples of temporal dynamics – of female choices and male traits – for the Cultural Model, the Gene-Culture Model, and their respective null models are shown in figure 3.

**Figure 3:**
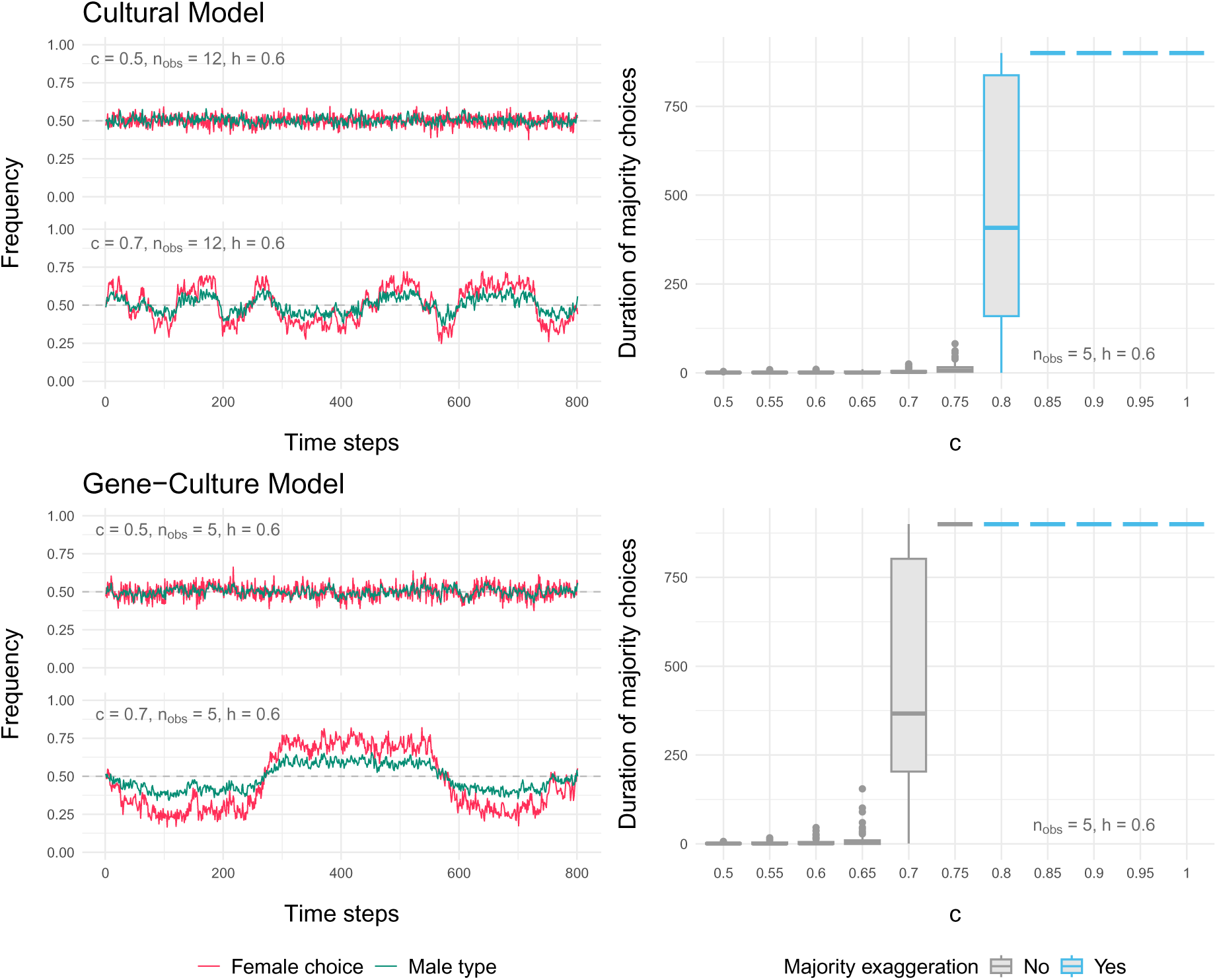
Examples of temporal dynamics and durations of majority behaviours. The first row corresponds to the Cultural Model and the second row to the Gene-Culture Model. On the left panel, temporal dynamics illustrate the evolution of female choice frequencies for type 1 males (red) and the frequency of type 1 males (green) over time. For both models, the top graphs represent the null model (*c* = 0.5) while the bottom graphs display the corresponding social models. The Cultural Model is shown for *n_obs_*= 12, *c* = 0.7 and *h* = 0.6, while the Gene-Culture Model is illustrated with *n_obs_* = 5, *c* = 0.7 and *h* = 0.6 (we chose different parameter sets in order to show interesting dynamics). On the right panel, boxplots represent the duration of majority choices, defined as the number of time steps during which the most frequent choice remains above 50%. Each boxplot summarizes results from 100 simulation replicates. Simulations were run with *n_obs_* = 5 and *h* = 0.6, while varying *c*. Blue boxplots correspond to conformity curves exhibiting majority exaggeration, whereas grey boxplots represent conformity curves without majority exaggeration.

In cases where one of the two male traits became permanently fixed (which can occur when *h* = 1), traditions could no longer be identified, as the majority choice would persist indefinitely regardless of social transmission. In our simulations, we therefore stopped measuring persistence duration once fixation occurred.

#### 2.3.2 Computing the proportion of traditions detected across simulations replicates

To identify the conditions that promote the emergence of traditions, we ran simulations while systematically varying the three key parameters: *n_obs_* and *c* (which determine the fidelity of social transmission) and *h* (which influences the fidelity of genetic transmission). All other parameters were held constant at their baseline values. Namely, population size (*K*) is fixed to 1000 individuals. The probability for juveniles to transition to adulthood at each time step (*a*) is set to 0.9, creating overlapping generations. This demographic structure prevents artificial cyclic dynamics that can occur with non-overlapping generations. The number of matings per male (*m*) is set to 100, which effectively allows for unlimited mating opportunities given the population size. The population is initially in a fully neutral state, with no pre-existing tradition: the initial average frequency of male trait 1 (*p*_0_) is set to 0.5 (i.e., an equal proportion of both male types), and *t*_0_ adult females have no preference, choosing mates at random (but are models for learning juvenile females). Each simulation was replicated 100 times, and figure 4 presents the proportion of simulations in which traditions were detected for each parameter combination.

**Figure 4:**
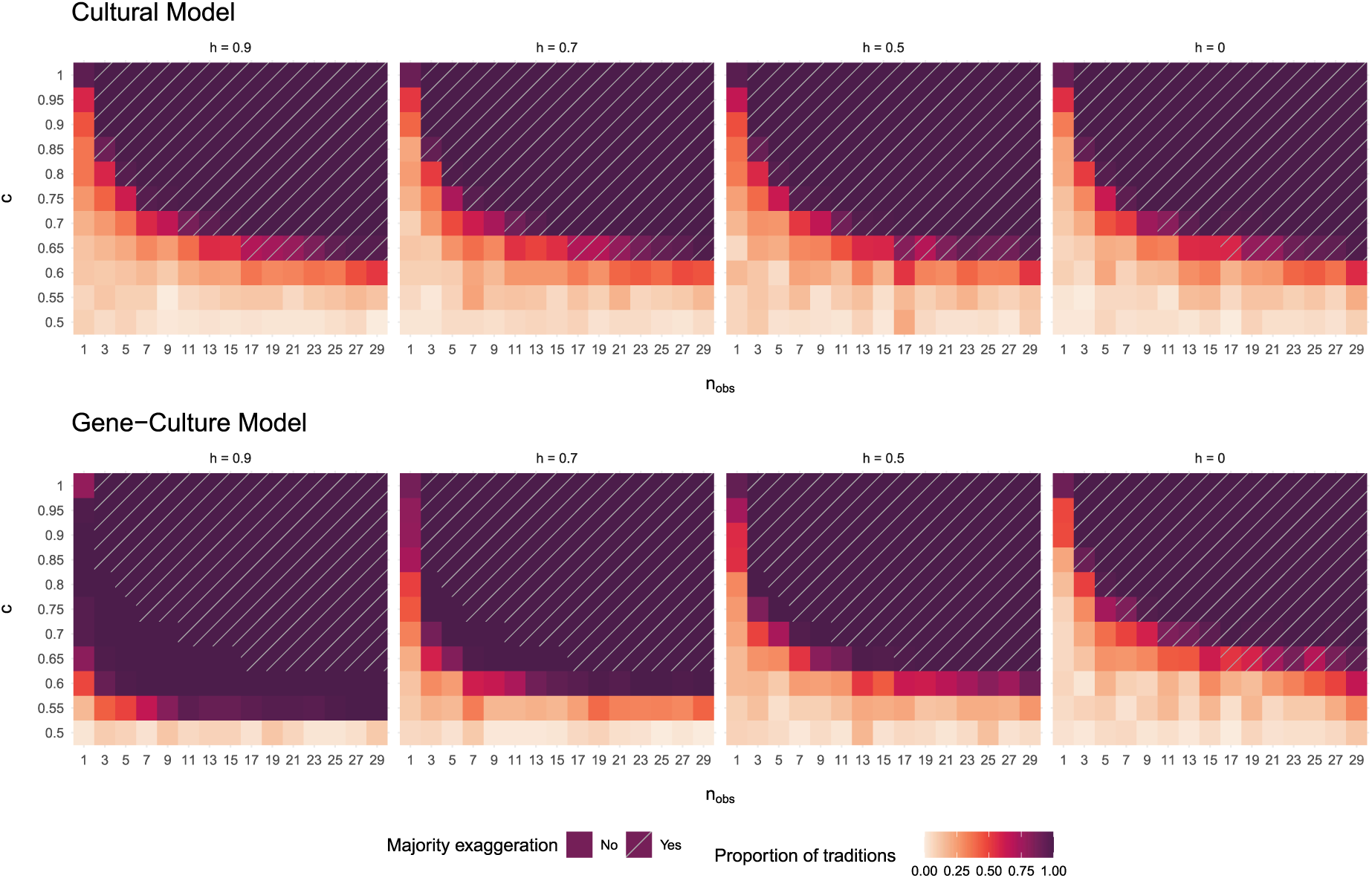
Proportion of traditions detected in simulations. The top row corresponds to the Cultural Model, and the bottow row to the Gene-Culture Model. Colours indicate the proportion of traditions detected across 100 simulation replicates, for different combinations of *n_obs_*, *c* and *h*. Traditions are defined as majority choices persisting beyond the detection threshold. This threshold is defined as the 95th percentile of majority durations observed in the matching null model – that is, the model with the same *n_obs_* and *h* but with *c* = 0.5. Grey-striped cells represent parameter combinations that induce majority exaggeration in the corresponding conformity curve.

Additional analyses are presented in the Supplementary Materials. These include robustness checks on our tradition detection method, with a test of the effect of the number of initial time steps discarded before measuring traditions (Section B, figure S2). The effects of varying initial conditions are explored in Section C (figure S3), by testing the influence of different initial frequencies of preferences and traits in the population. This analysis also serves as a preliminary investigation of the relative strength of gene–culture interactions in our models. The impact of limiting the number of matings per male (*m*) is explored in Section D (figure S4), allowing to explore alternative mating systems (different strength of polygyny as well as monogamy). The consequences of varying demographic parameters (*K* and *a*) are examined in Section E (respectively in figures S5 and S6).

#### 2.3.3 Traditions in the Cultural Model

As expected, the proportion of simulation replicates in which a tradition is detected increases with both *n_obs_*and *c* (figure 4, first row). More specifically, in the Cultural Model, traditions cannot emerge unless the conformity curve exhibits majority exaggeration. When *n_obs_* and *c* are too low to produce this effect, the majority behaviour is not reinforced, preventing the formation of traditions. Conversely, as soon as the conformity curve exaggerates the majority behaviour, even slightly (dashed area in figure 4, first row), traditions emerge.

Notably, in the Cultural Model, genetic transmission fidelity (*h*) has no impact on the emergence of traditions. This is because, by construction, as long as both male types are available, female mate choice remains independent of male type frequencies (see figure 1). In other terms, the dynamics of mate choice in this model is not influenced by the genetic evolution of male traits.

#### 2.3.4 Traditions in the Gene-Culture Model

In the Gene-Culture Model, we observe the same general trend as in the Cultural Model: the higher *n_obs_* and *c*, the more likely traditions are to emerge (figure 4). However, a key difference is that, in the Gene-Culture Model, traditions can arise even in the absence of majority exaggeration in the conformity curve (see grey boxes in figure 3 and non-dashed areas in figure 4, second row).

This is because, in the Gene-Culture Model, conformist learning is not the only mechanism reinforcing majority behaviours. Since female mate choice also depends on male type availability, an interaction emerges between female choice and male trait frequencies. Once a trait becomes predominant due to stochastic fluctuations (be they demographic fluctuations or fluctuations in mate choice), it will be encountered and chosen by adult females, which in turn will make juvenile females prefer it. The preference will in turn increase the frequency of that trait in sons and strengthens the preference for it in daughters. This self-reinforcing cycle continues, until choice and trait frequencies stabilize at an equilibrium determined by a non-trivial balance between social and genetic transmission fidelity.

The strength of this frequency-dependent effect is directly influenced by the fidelity of genetic transmission: the higher *h*, the stronger the reinforcement between choice and trait, and the lower the conformist capacities needed to sustain traditions. For instance, with *h* = 0.9, traditions emerge in 100% of simulation replicates with even moderate conformist capacities (for instance, *n_obs_* = 5 and *c* = 0.65), and even single-copying (*n_obs_* = 1) can generate long-lasting traditions if *c ≥* 0.65.

Conversely, when *h* = 0, male traits are not heritable, which prevents the self-reinforcing cycle effect. In this case, the Gene-Culture Model converges to the Cultural Model, meaning traditions can no longer emerge without majority exaggeration in social learning.

## 3 Discussion

### 3.1 Summary of the study’s goals

This theoretical study investigates the conditions under which traditions emerge in mate choice, using simulation models where female mate preferences are socially transmitted through conformity, while preferred male traits are genetically inherited. We examined how traditions become established under different levels of social transmission fidelity by varying two key parameters influencing the intensity of conformist learning: the number of observations *n_obs_* and the copying fidelity *c*. In particular, we investigated whether majority exaggeration is a necessary feature of conformity for traditions to emerge. We first addressed this question in a model where mate choice is not constrained by the frequency of male types in the population (Cultural Model), allowing the cultural evolution of female preferences to occur independently of the genetic evolution of male traits. We then studied a model in which female choice is constrained by male-type frequency, creating a gene–culture coevolutionary context – specifically, a coevolution between male traits and female preferences (the Gene–Culture Model).

### 3.2 Traditions without genetic influences in the cultural model

In the Cultural Model, mate choice is ‘unconstrained,’ meaning that a female’s choice is determined solely by her level of conformist learning – measured by parameters *n_obs_*and *c* – regardless of the frequency of her preferred male type in the population (see figure 1, in orange). Our results show that a wide range of *n_obs_* and *c* combinations lead to the emergence of traditions (figure 4). Consistent with previous studies, we find that traditions hardly emerge without majority exaggeration in the conformist response curve [36, 18, 39] (figure 4). Conformist learning must produce majority exaggeration for traditions to arise in this model, as this is the only mechanism that creates a form of positive frequency-dependent selection, promoting the persistence of any behaviour that temporarily becomes the majority due to random fluctuations. This model reflects the standard approach in cultural evolution studies, where social transmission is the primary evolutionary force and where individuals’ choices are not constrained by resource availability – in other words, where preferring and choosing a behaviour are not clearly distinct (e.g., when individuals choose between two or more techniques to open a box, to wash food, or when they decide which song to sing, etc.). This allowed us to establish baseline results for our modelling of conformity with *n_obs_*and *c* before introducing a gene–culture coevolutionary context.

### 3.3 Traditions are favoured by a gene-culture coevolution context

We show that traditions emerge most readily in a model that is arguably more realistic, where mate choice is constrained not only by females’ conformist abilities, but also by their opportunities to encounter preferred male types. As a result, in this Gene–Culture Model, female choice partly reflects the frequency of male traits in the population, creating a context of gene–culture coevolution (see figure 1, in blue). When male traits are heritable (*h >* 0), this scenario generates a positive feedback loop that reinforces the persistence of majority behaviours, thereby promoting the establishment of traditions (figure 4).

Introducing the mate encounter step in the Gene-Culture Model amounts to making a clearer distinction between the concepts of female ‘preference’ and ‘choice’. Traditions result from choices, which depend in this model not only on preference but also on the availability of the resource (i.e., the male type). This encounter dynamic can be seen as a repair mechanism for traditions: females who misestimate the population majority choice and develop a preference for the minority type still have a high chance of mating with the majority type – which correlates with the majority choice – simply because they encounter it more frequently. This mechanism effectively counteracts copying errors produced by imperfect conformity, thereby facilitating the establishment of traditions. The distinction between preference and choice is common in the context of mate selection [51, 52] (and is sometimes reflected in the concepts of preference and choosiness [42]). However, in classic models of conformity, to prefer a behaviour is assumed to mean adopting it. We argue that this equivalence should be reconsidered for all culturally transmitted behaviours that depend on access to a resource, as our results show that resource dynamics can significantly alter – and in this case, substantially facilitate – the conditions under which traditions emerge.

Previous sexual selection models have described positive frequency-dependant dynamics between genetically inherited traits and preferences acquired by social learning [43, 44, 45] or by sexual imprinting [53, 54]. While related to this family of models, our study differs by focusing on cultural evolution and traditions rather than on genetic consequences, and by explicitly modelling conformist transmission (though see [55] for conformity effects on male morph dynamics). Notably, in Duval et al.’s study [56], the importance of social preferences for traditions is acknowledged, but through a social learning mechanism in which preferences are shaped by estimating the rarity of traits in potential mates – in contrast to our study, where social preferences are based on majority detection.

The model most closely related to ours is that of Kirkpatrick and Dugatkin [43]. Using a population genetics framework, they explored two mate-copying rules: ‘single copying’, where females copy the choice of a single older female, and ‘mass copying’, where they copy the majority choice among all older females – rules that correspond to *n_obs_* = 1 and *n_obs_* = *M* 0 + *M* 1 in our study. Both rules produced positive frequency-dependent dynamics, and the authors pro-posed as a general mechanism that copying (as well as sexual imprinting) reinforces female preferences for already-preferred male types. Building on our results, we suggest that this interpretation holds only if copying fidelity is sufficiently high and, crucially, if copying occurs in a context that allows preferences and traits to coevolve. This coevolution emerges in their models – just as in our Gene-Culture Model – because female choices depends not only on preferences but also on the availability of preferred males. Kirkpatrick and Dugatkin [43] do not explicitly discuss this point, but the influence of mate availability is implicit in their equations 3 and 4. Once again, we emphasize the importance of distinguishing preference from choice in future studies to better understand the respective roles of social learning and other interacting factors in shaping cultural dynamics.

Kirkpatrick and Dugatkin [43] questioned the relevance of comparing the positive frequency-dependent dynamics emerging from their models to classical reinforcement processes in sexual selection theory, such as the Fisher runaway process [40]. They noted that some of the similarities between their models and classical models are merely superficial, since, for example, the reinforcement dynamics in their models cannot be attributed to linkage disequilibrium between preferences and traits. The same point applies to our models. We could add that, unlike the original Fisherian model – which was proposed to account for the evolution of arbitrarily exaggerated secondary sexual traits (such as the costly peacock tail) and corresponding preferences [40] – our models focus exclusively on the frequencies of traits and preferences. In other words, in our framework, a trait becomes ‘exaggerated’ not in terms of its phenotypic expression or cost, but simply by being more frequently chosen by females. Despite these differences, we argue that the underlying evolutionary mechanisms show strong parallels. As in Fisher’s original model, coevolution in our study arises from a female preference that can be described as arbitrary. Here, ‘arbitrary’ means that conformist females tend to prefer whichever male trait initially deviates randomly from a frequency of 1/2 – whereas in the Fisher model, they tend to prefer whichever trait has a value that deviates from the mean. Another putative strong parallel is that if our model were modified to include the possibility for copying mechanisms to evolve, our modified model would produce a version of the ‘sexy son’ effect: the indirect selective advantage for females who correctly copy the majority would lie in producing sons who carry the traits most likely to be attractive to the next generation of females – as well as daughters most likely to prefer the most attractive males at the next generation. In this study, we show that this sexual selection on males can lead to the emergence of traditions for either of two initially neutral traits, and based on previous studies [43, 44], we expect it may also be sufficient to generate traditions for a costly trait.

Overall, our results reinforce the idea that mate choice, though often overlooked, is a particularly relevant context for studying cultural evolution patterns [28]. Sexual selection is one of the most extensively studied examples of gene-gene coevolutionary dynamics, but gene-culture coevolution may also be widespread in sexual selection and across many other biological scenarios [25, 57], exerting similar reinforcing effects on traditions. After being well-documented for human evolution (see for instance [58]), gene–culture coevolutionary processes are increasingly being identified in non-human species [27], raising the question of how such interacting evolutionary processes might influence traditions.

More broadly, this perspective aligns with the growing recognition that interactions between genetic and non-genetic inheritance systems must be more fully integrated into our understanding of evolutionary processes. Our findings also align with a growing body of work advocating for a broader perspective on cultural evolution – one that extends beyond social transmission alone [21]. Recent studies have emphasized the importance of considering how social transmission mechanisms interact with broader population-level factors [24]. For instance, research has begun to examine how social learning interacts with variables such as population size [18], age structure [22], social network structure [23], and migration [19]. As demonstrated in our study with the example of gene-culture coevolution, integrating these broader perspectives can refine conclusions about the mechanisms underlying cultural patterns.

### 3.4 Majority exaggeration is not always needed for traditions to emerge

In cultural evolution research, conformity is often defined – following Boyd and Richerson’s original proposal – as the disproportionate probability of adopting the majority behaviour [36, 46, 38, 28, 59, 39], a feature we refer to as ‘majority exaggeration’ (figure 2). In this study, we adopt a more mechanistic approach to modelling conformity. Rather than imposing a predefined shape for the population-level conformist response, we modelled individuals who engage in a conformist learning process, and derived conformist curves from the models. Some of the emerging conformity curves display majority exaggeration under certain parameter sets, but not all (figure 2). In other words, we examined conformity both in the presence and absence of majority exaggeration. Our approach is similar to that of Claidière and Whiten [60, 48], who propose to classify different types of conformity based on their effects on behavioural acquisition curves, distinguishing ‘hyperconformity’ (which involves majority exaggeration) and ‘weak conformity’ (which does not).

Contrary to common predictions [39], we found that traditions can emerge even without majority exaggeration in conformist learning (figures 3 and 4, Gene-Culture Model). This is possible because, in a coevolutionary context, as discussed above, conformity is no longer the sole mechanism reinforcing majority behaviours. As a result, even minimal levels of conformist learning (e.g., as low as *n_obs_*= 3 and *c* = 0.6 for *h* = 0.9, figure 4, Gene-Culture Model) are sufficient to establish long-lasting traditions when traits and choices coevolve. Surprisingly, even single-copying (*n_obs_* = 1) – where individuals copy a randomly encountered conspecific without actively sampling or detecting majorities – can give rise to highly lasting traditions. This finding contradicts previous results claiming that ‘one cultural parent makes no culture’ [61] and may be particularly relevant for future research on the consequences of mate-choice copying, which has largely focused on single-copying in experimental settings. Overall, our results suggest that cultural evolution can emerge from socio-cognitive abilities that are less advanced than commonly assumed [39, 61], at least in mate choice.

In most previous studies, conformity has been modelled using mathematical functions that generate sigmoidal curves, with *ad hoc* parameters controlling, for instance, the slope at the inflection point [36, 62, 39]. These parameters are set to represent the strength of conformity, but they are rarely given a clear biological interpretation. Moreover, the same sigmoidal curve could result from different mechanisms at the individual level [47]. By modelling conformity at the individual level using the parameters *n_obs_* (number of observations) and *c* (fidelity of copying), we captured diverse ways in which individuals exhibit conformist behaviour. These parameters may reflect cognitive abilities and/or learning strategies, encompassing various ways individuals sample, discriminate quantities to sense a majority, memorize, or decide to rely on social information. Overall, we suggest that an individual-level approach to modelling conformity can help identify biologically meaningful parameters – making them more interpretable than in previous studies and more directly comparable to measurable variables in experimental work. Redefining conformity in terms of individual learning mechanisms, without restricting the shape of conformity curves to the ones with majority exaggeration, could help minimize assumptions about evolutionary outcomes and allow for a broader exploration of social learning strategies based on majority detection. This, in turn, could improve predictions about which species and ecological contexts are most likely to support traditions.

The robustness of our main findings was systematically evaluated across varying demographic parameters, mating systems, and initial conditions (see Supplementary Materials, Sections C-E). Our results remain robust across demographic variation, confirming that the emergence of traditions through gene-culture coevolution is not contingent on specific demographic assumptions (Section E). These analyses also reveal important nuances: for example, we show that strict monogamy substantially constrains tradition emergence, requiring higher genetic and social transmission fidelity compared to polygynous systems (Section D). These nuances call for comparative studies across species with varying mating systems and modes of inheritance to better understand the ecological and evolutionary conditions that promote cultural traditions. Furthermore, analyses of initial conditions indicate that in the Gene-Culture Model, the genetic component (trait frequency) has a stronger influence than the social component (preference frequency) in determining the direction of emerging traditions (Section C). In this study, we have adopted the standard definition that traditions must result, at least in part, from social transmission. However, this definition does not specify how important social transmission must be relative to other mechanisms. If the genetic component has greater influence than the social component in establishing traditions, can we still reasonably speak of traditions? It may be useful to view culture as a gradual property – some traditions being more or less cultural depending on the relative contributions of social and non-social factors [63, 64]. Traditions are not always ‘purely social’, and one challenge for future research will be to better understand which factors promote traditions and which constrain them.

### 3.5 Perspectives

This study leaves several aspects open for future work. We considered a scenario with binary traits and preferences, but as suggested by previous studies, traditions may be more difficult to establish in a potentially more realistic context where conformity applies to more than two traits [50], or to continuous traits. The mechanisms of social transmission in our models could also be refined to better reflect biological realism. Empirical evidence suggests that preference acquisition is often highly flexible and varies within species, a factor not currently captured in our models. Future studies could account for the fact that individuals may rely on social or personal information to varying degrees depending on the context [65, 66, 67, 68], and that there may be inter-individual variation in social learning related to stable personality traits [69, 70, 71, 72]. Finally, although we assumed a theoretical mating system in which females are the choosy sex, many species that are promising candidates for cultural traditions exhibit some degree of mutual mate choice — as observed in certain passerine birds, primates, and even flies [73]. Finally, we modelled conformity as the default and established strategy for mate choice, allowing us to focus on the levels of conformity that lead to the emergence of traditions. In future work, it will be interesting to examine the evolution of learning parameters and the conditions under which conformity itself evolves as an evolutionarily stable strategy – an emerging topic in the study of mate choice [28]. Understanding the evolution of conformity will be a key step in refining our knowledge of the conditions required for traditions to persist in mate choice.

## 4 Data, Scripts and Code Availability

Scripts of the models, data from simulations and scripts of data analysis can be found on Zenodo at https://doi.org/10.5281/zenodo.17144302 [74].

## 7 Acknowledgements

We thank Manuel Sapage (reviewer), one anonymous reviewer and Alexandre Courtiol (recommender) for their insightful advice and valuable feedback on our study. We also thank Eric Lombaert and Alexandra Auguste (data editors for PCI), who helped us improve the reproducibility and reusability of our scripts and simulation data. We are grateful to Marlène Framit, Hugo Ostrowiecki and Mattis Gautrand for their contributions during research internships on topics related to this study.

## Supplementary materials

### A Conformity curves: comparison between analytical and simulations curves

We defined conformist curves in the main text of the article as the relationship between the probability that a female chooses a male of type 1, conditional on encountering one, *P* (*C*_1_ *| E*_1_) at time *t*, and the frequency of choices for type 1 in the population, *F* (*C*_1_), at time *t −* 1. In the main text, we develop an analytical method to calculate *P* (*C*_1_ *| E*_1_) (see equations 1 and 2). The aim of this section is to estimate *P* (*C*_1_ *| E*_1_) from simulation data in order to verify that our models produce the conformist curves predicted by our analytical method. A comparison between curves from simulations and their analytical counterparts is shown in figure S1.

In our Cultural Model, *P* (*C*_1_ *| E*_1_) = *P* (*C*_1_) (as detailed in the main text), and *P* (*C*_1_) can be estimated by computing *F* (*C*_1_), the frequency of choices for type 1. Thus, for the Cultural Model, the conformist curves derived from simulations are estimated from the relationship between *F* (*C*_1_) at time *t*_1_ and *F* (*C*_1_) at time *t*_0_ (see figure S1A). In our Gene–Culture Model, the probability that a female chooses a male of type 1 given that she has encountered one is difficult to measure directly from simulations, but can be reasonably approximated from the probability of preferring a type 1 male, *P* (*Pref*_1_), which is easily obtained from simulation data. Hence, *P* (*C*_1_ *| E*_1_) = *cP* (*Pref*_1_) + (1 *− c*)(1 *− P* (*Pref*_1_)). Since *P* (*Pref*_1_) can be estimated as *F* (*Pref*_1_), the conformist curves from simulations are given by the relationship between *cF* (*Pref*_1_) + (1 *− c*)(1 *− F* (*Pref*_1_)) at time *t*_1_ and *F* (*C*_1_) at time *t*_0_ (see figure S1B). Curves from simulations correspond to mean curves calculated from 1,000 simulation replicates, and the parameters used in these simulations are fixed to their baseline values (as described in the main text).

**figure S1:**
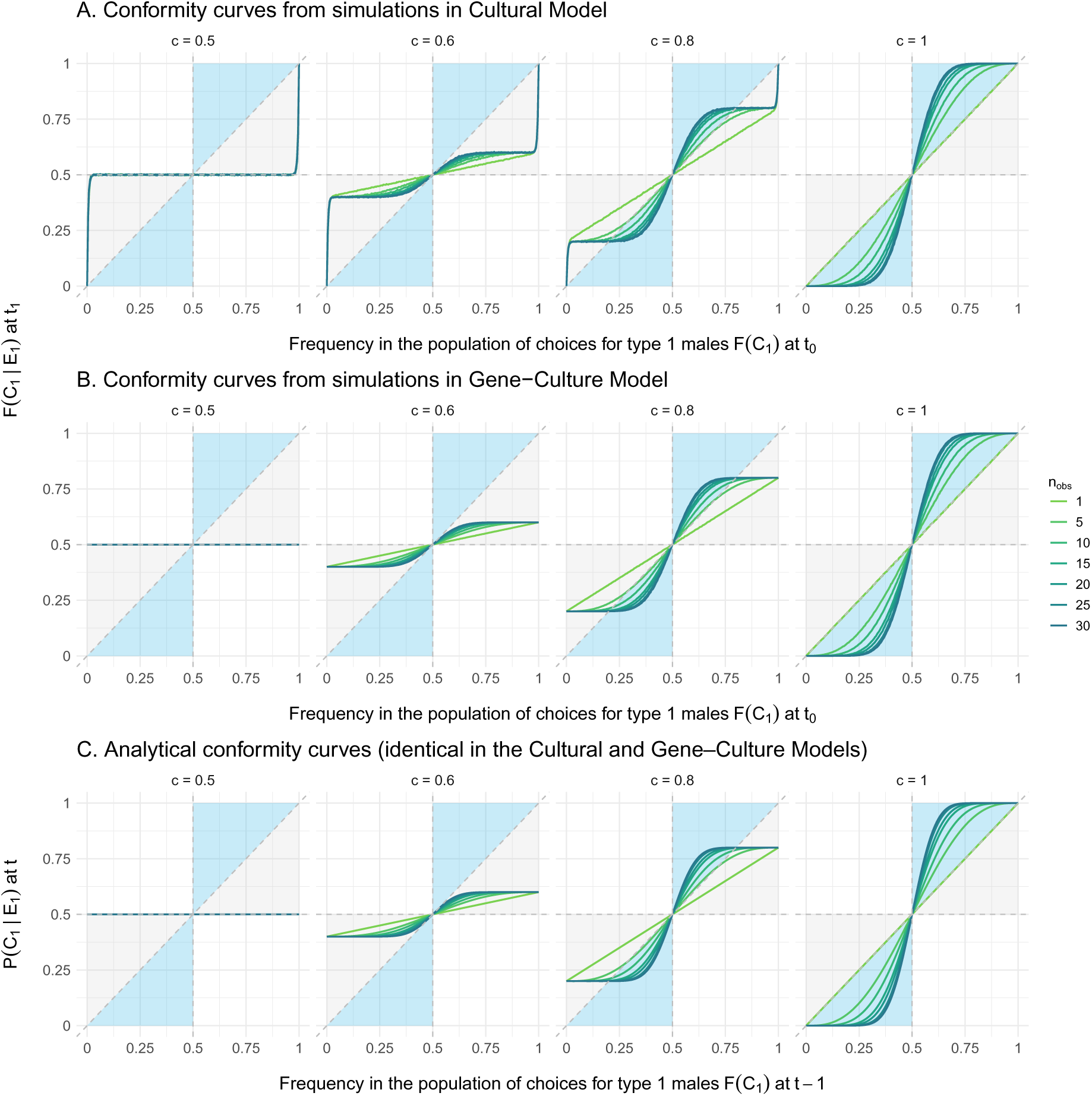
Conformity curves from simulations (A and B) versus analytical conformity curves (C). The curves derived from simulations closely match the analytical predictions (figure S1), except when *F* (*C*_1_) approaches 0 or 1 (visible for the Cultural Model and expected as well for the Gene–Culture Model, though not visible with our current estimation of *P* (*C*_1_ *| E*_1_)). This discrepancy arises because, in our simulations, once a trait is fixed in the population, choices for the alternative trait become impossible — a situation not accounted for in the analytical method. However, this edge case never occurs in the parameter settings presented in the main text results.

### B Number of time steps removed before starting the detection of traditions

In the method used to detect traditions in the main text, we excluded the first 100 time steps of each simulation before beginning the measurement of traditions, in order to avoid potential effects of initial conditions on the dynamics. Since the number of excluded time steps is somewhat arbitrary, we here test the robustness of our results to variations in this number. Specifically, we compute the proportion of traditions detected across 100 replicates for each simulation (using the same method as in the main text), after removing either 25, 50, 100, or 200 initial time steps, for both the Cultural Model and the Gene–Culture Model (figure S2). In these analyses, we vary the parameters *n_obs_* and *c* as in the main text, and fix the genetic transmission fidelity at *h* = 0.9, which corresponds to a scenario in which gene–culture interactions can promote the emergence of traditions in the Gene–Culture Model. All other parameters are held at their default values (i.e., those used in the main text). For the tested parameter values, we find that our results on the emergence of traditions are robust to changes in the number of time steps removed prior to measuring traditions (see figure S2).

**figure S2:**
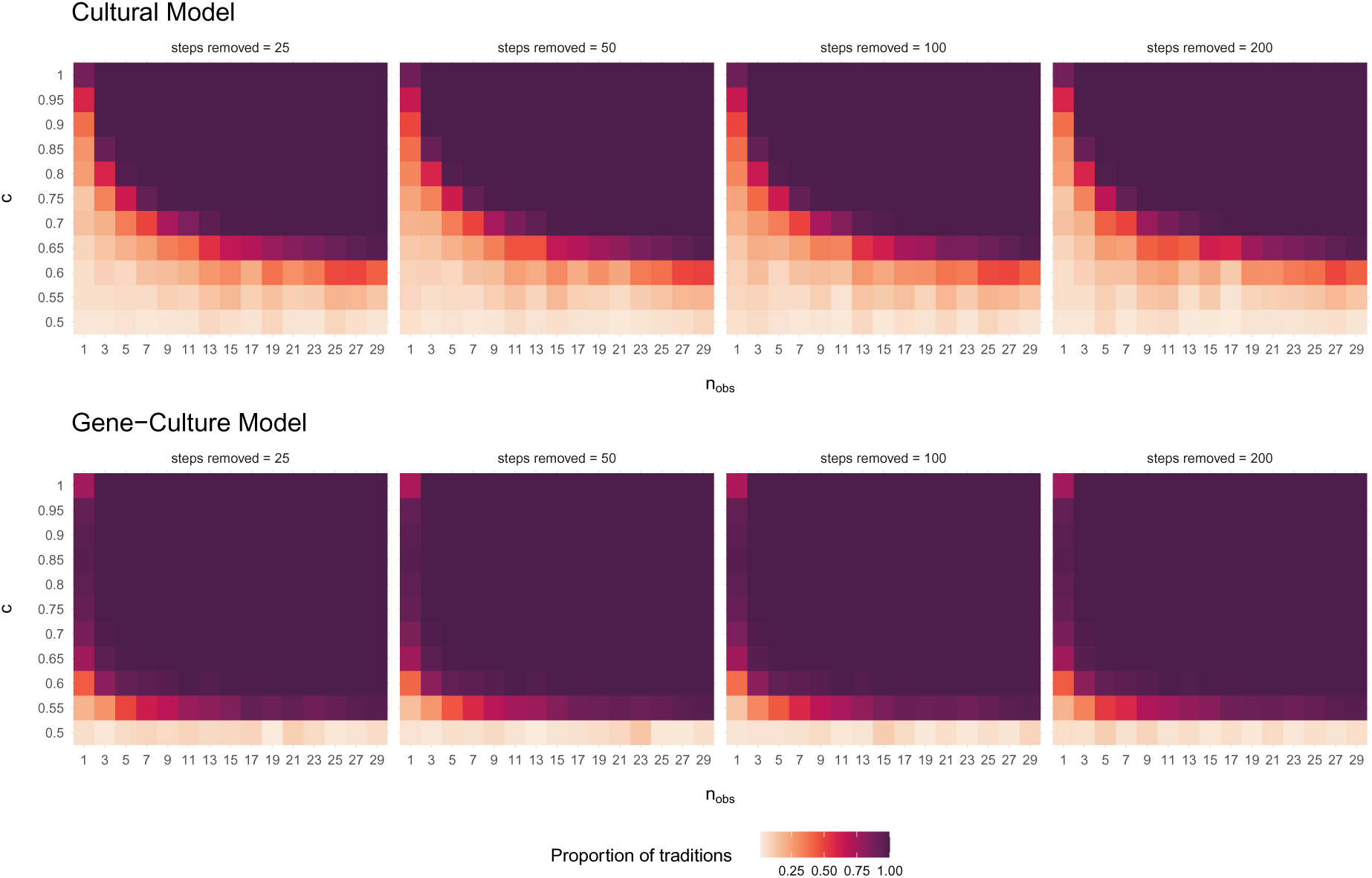
Effect of the number of time steps removed in the method used to detect traditions on results about traditions. The colour corresponds to the proportion of traditions detected in 100 simulation replicates, for different values of the parameters *n_obs_*, *c* and number of time steps removed, for the Cultural Model and the Gene-Culture Model.

### C Initial conditions: a measure of gene-culture interactions

The results presented in the main text correspond to neutral initial conditions: at time step *t*_0_, the initial frequency of male trait 1 is on average 0.5; adult females at *t*_0_ have no preference and choose mates at random; and the average preference of juvenile females at *t*_1_ is also 0.5, since they socially acquire their preference by observing the random choices made by adult females at *t*_0_. These neutral initial conditions allowed us to isolate the effects of conformist learning and gene–culture coevolution on the establishment of traditions — and to study the emergence of traditions from so-called arbitrary preferences.

In the analysis below (figure S3), we simultaneously vary the initial frequency of trait 1 in males and the initial frequency of preference for trait 1 in females, and we measure the impact of these starting conditions on the outcome of the first emerging tradition (i.e., whether the first tradition is for trait 1 or trait 0). Specifically, we compute the proportion of first traditions for trait 1, based on 100 replicates per simulation setting (parameter values used in the simulations are provided in the legend of figure S3). This analysis allows us to test the influence of different initial configurations regarding the prevalence of preferences and traits in the population. It also serves as a preliminary investigation of the relative strength of gene–culture interactions in our models. In particular, it helps determine whether the social component (via initial preference frequency) or the genetic component (via initial trait frequency) has the greater effect on the direction of the first tradition.

In the Cultural Model, it is the social component (initial preference frequency) that fully determines the direction of the first tradition (figure S3, top row). This result is expected, since by design, the frequency of male genetic traits does not influence female choice in this model. In contrast, in the Gene–Culture Model, it is the initial frequency of the trait (the genetic component) that appears to have the strongest influence on the direction of the first tradition (figure S3, bottom row), while the initial frequency of the preference (the social component) seems to have little effect. Of course, the emergence of traditions still necessarily relies, at least in part, on social transmission — since without it, gene–culture coevolution could not occur. However, this result highlights the major role of the genetic component in the establishment of traditions in our gene–culture framework.

**figure S3:**
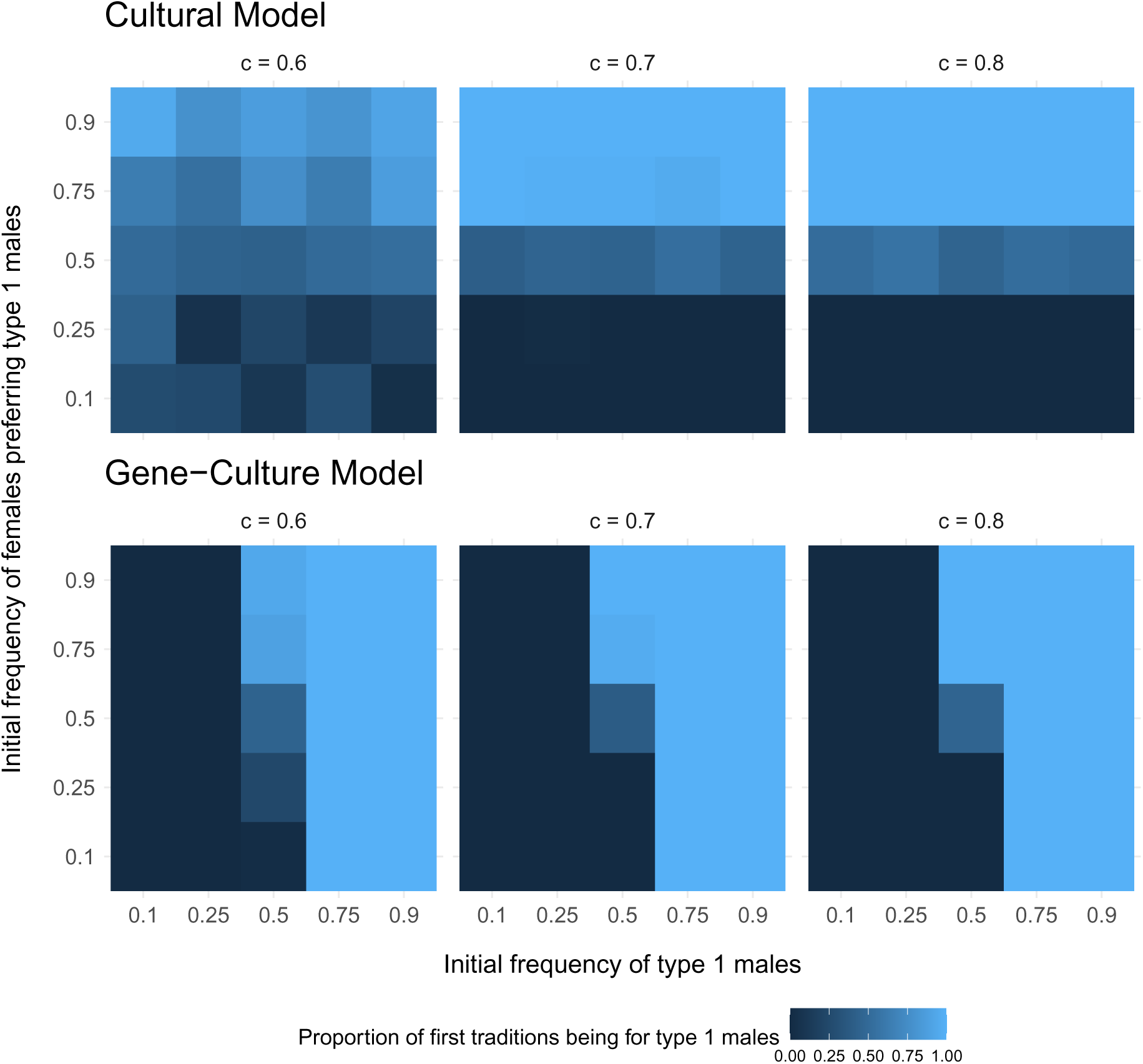
A measure of gene–culture interactions: effect of initial conditions on the first tradition generated by conformist learning. Colour indicates the proportion of first traditions corresponding to a majority choice for type 1 males (a “tradition for type 1 males”). The first tradition is defined as the first majority preference that arises during the simulation and whose duration is significantly longer than those observed in the corresponding null model. This proportion was computed from 100 simulation replicates for: (1) varying initial frequencies of females preferring type 1 males (y-axis), (2) varying initial frequencies of type 1 males in the population (x-axis), (3) different values of the copying fidelity *c* (separate heatmaps), and for the two models, Cultural Model (top row) and Gene–Culture Model (bottom row). The number of observations and the genetic transmission fidelity were fixed at *n_obs_*= 7 and *h* = 0.95, respectively.

### D Restricting the number of matings per male: from polygynous to monogamous species

In our Gene–Culture Model, the parameter *m* determines the maximum number of matings an adult male can achieve. The results presented in the main text are based on a polygynous species, in which males can potentially mate with multiple females, with this maximum set to *m* = 100 — a value that is never reached in practice in populations of 1,000 individuals.

In the analysis below (figure S4), we test the effect of the number of allowed matings per male on the emergence of traditions. figure S4 shows the proportion of traditions obtained over 100 simulation replicates, for various values of the maximum number of male matings (the simulation parameters are detailed in the legend of figure S4). This analysis allows us to: (1) test the robustness of our results to changes in this parameter, and (2) explore alternative mating systems compared to our initial model. By lowering the number of matings, we can for instance simulate a monogamous species (in which each individual mates only once).

In scenarios that lead to long-lasting traditions in the initial model — for instance, with *h* = 0.95, or *h* = 0.8 and *c ≥* 0.7 — we observe a threshold effect regarding the impact of available male matings (figure S4). For *m ≥* 2, traditions are unaffected or only minimally affected by male mating constraints. Only the transition to a strictly monogamous species (*m* = 1) markedly limits the establishment of traditions. While traditions can still emerge under monogamy, they require very high fidelity in both genetic and social transmission.

**figure S4:**
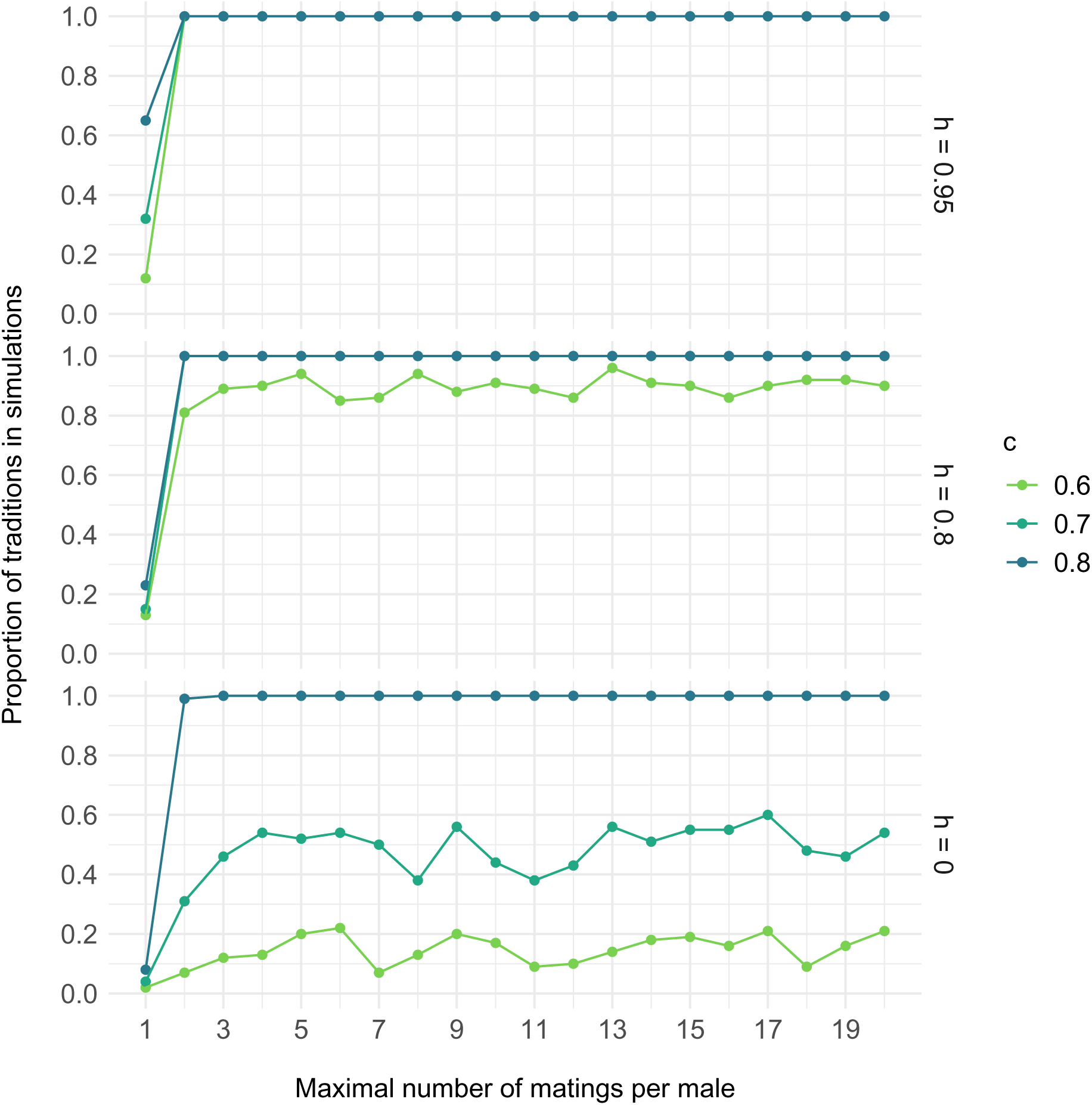
Effect of the maximal number of matings per male on the emergence of traditions through conformity. The proportion of traditions is computed from 100 replicates per simulation. The number of observations is fixed at *n_obs_* = 7. Results are shown for selected values of the parameters *c* and *h*, to illustrate scenarios with either high or low levels of tradition emergence. Simulations were conducted using the Gene–Culture Model.

### E Robustness to demographic parameters

Finally, we tested the robustness of our results on the emergence of traditions to variation in the model’s demographic parameters – namely, population size (*K*) and ageing rate (*a*). The results presented in the main text correspond to the following default values: *K* = 1000 and *a* = 0.9. Here, we compute the proportion of detected traditions over 100 simulation replicates (using the same method as in the main text), across different values of *K* (figure S5) and *a* (figure S6), for both the Cultural Model and the Gene–Culture Model. In these analyses, we vary the parameters *n_obs_* and *c* as in the main text, and set the genetic transmission fidelity to *h* = 0.9, which corresponds to a scenario where gene–culture interactions can promote the emergence of traditions in the Gene–Culture Model. Parameters not being tested are kept at their baseline values (i.e., those used in the main results). For all tested parameter values, we find that our results on tradition emergence are robust to variation in *K* and *a* (see figures S5 and S6).

**figure S5:**
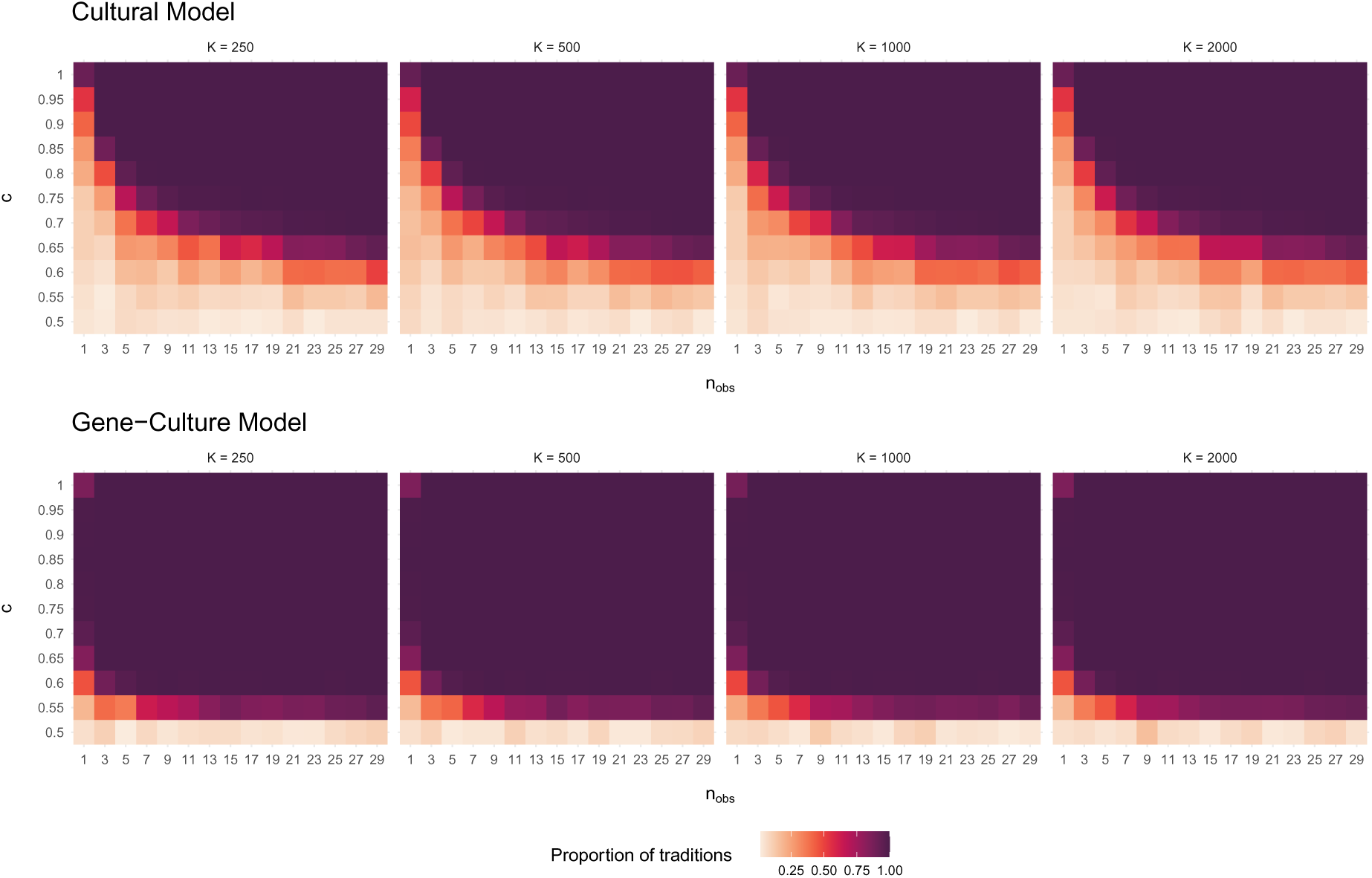
Effect of parameter *K* (population size) on traditions.

**figure S6:**
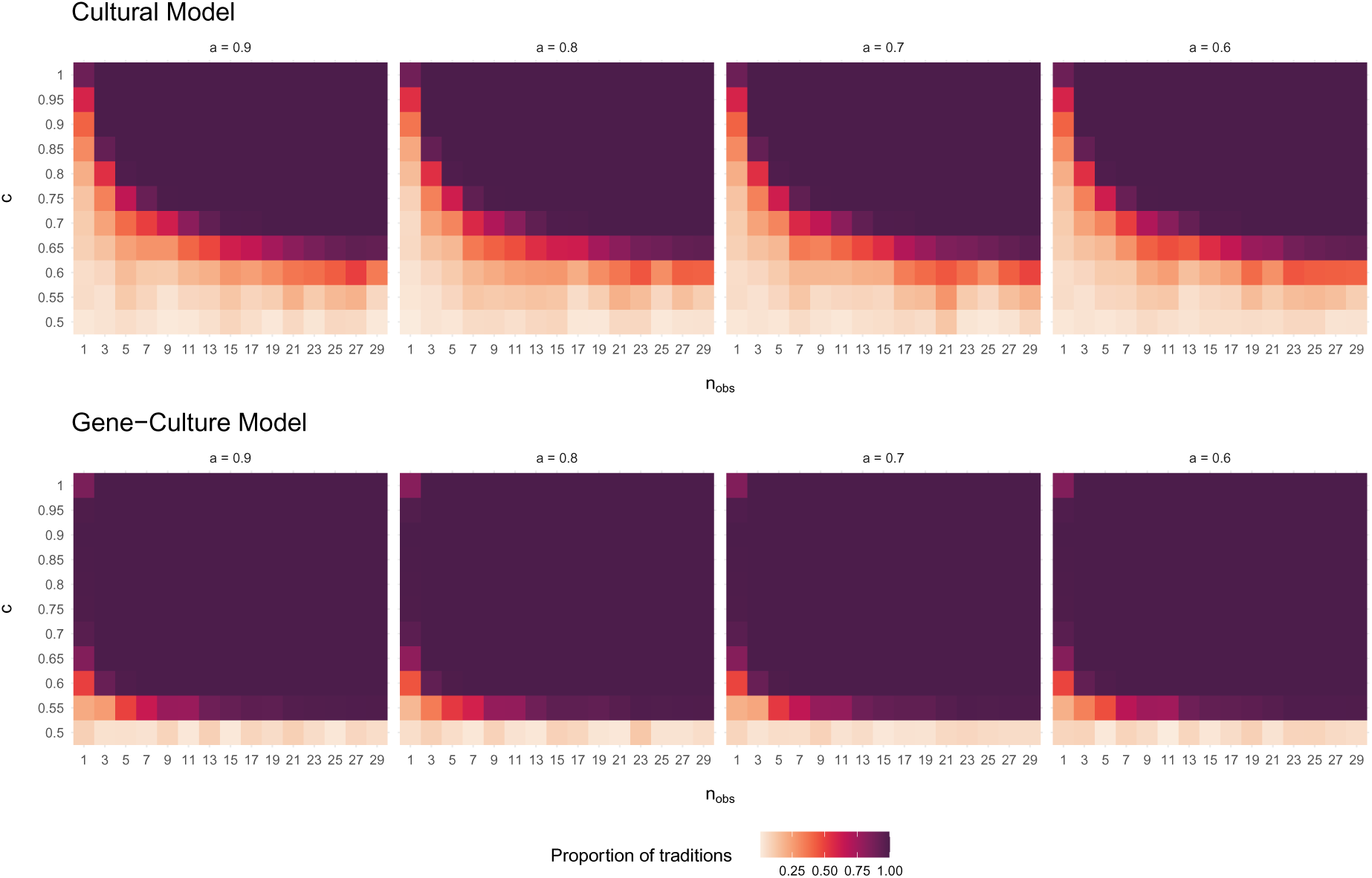
Effect of parameter *a* (ageing rate) on traditions.

